# Evaluating intra- and inter-individual variation in the human placental transcriptome

**DOI:** 10.1101/012468

**Authors:** David A Hughes, Martin Kircher, Zhisong He, Song Guo, Genevieve L. Fairbrother, Carlos S. Moreno, Philipp Khaitovich, Mark Stoneking

**Affiliations:** Max-Planck Institute for Evolutionary Anthropology, Deutscher Platz 6, Leipzig Germany 04103; CAS-MPG Partner Institute for Computational Biology, 320 Yue Yang Road, Shanghai 200031, P.R. China; Obstetrics and Gynecology of Atlanta, 1100 Johnson Ferry Rd NE Suite 800, Center 2, Atlanta, GA 30342, USA; Department of Pathology and Laboratory Medicine and Department of Biomedical Informatics, Emory University, Atlanta, GA 30322, USA

**Keywords:** transcriptome, gene expression, selection, apportionment, placenta, Nst, Mst, population genetics

## Abstract

**Background:** Gene expression variation is a phenotypic trait of particular interest as it represents the initial link between genotype and other phenotypes. Analyzing how such variation apportions among and within groups allows for the evaluation of how genetic and environmental factors influence such traits. It also provides opportunities to identify genes and pathways that may have been influenced by non-neutral processes. Here we use a population genetics framework and next generation sequencing to evaluate how gene expression variation is apportioned among four human groups in a natural biological tissue, the placenta.

**Results:** We estimate that on average, 33.2%, 58.9% and 7.8% of the placental transcriptome is explained by variation within individuals, among individuals and among human groups, respectively. Additionally, when technical and biological traits are included in models of gene expression they account for roughly 2% of total gene expression variation. Notably, the variation that is significantly different among groups is enriched in biological pathways associated with immune response, cell signaling and metabolism. Many biological traits demonstrated correlated changes in expression in numerous pathways of potential interest to clinicians and evolutionary biologists. Finally, we estimate that the majority of the human placental transcriptome (65% of expressed genes) exhibits expression profiles consistent with neutrality; the remainder are consistent with stabilizing selection (26%), directional selection (4.9%), or diversifying selection (4.8%).

**Conclusion:** We apportion placental gene expression variation into individual, population and biological trait factors and identify how each influence the transcriptome. Additionally, we advance methods to associate expression profiles with different forms of selection.

## Background

Nearly four decades ago, it was estimated that about 85% of the neutral genetic variation in humans is found within groups and only about 15% between groups [1], which reflects the close genetic relationship of human populations. This initial observation, using protein markers, has been substantiated by numerous additional studies and markers [2-6]. Further, these analyses provide a framework to identify genes that exhibit unusually large differences between populations and thus may have been subject to recent local positive selection [2, 7-10] as responses to population-specific evolutionary forces.

In principle, the variation in phenotypic traits can also be apportioned into within-population and between-population components [11], which could provide insights into the relative influence of both genetic and environmental factors on such traits. However, this has been done for only a few human traits. For example, cranial variation among human populations present between-population components (0.11 – 0.14) similar to neutral genetic variation [12], suggesting that human cranial variation also (largely) reflects neutral genetic processes. Conversely, variation in skin pigmentation has a significantly larger between-population component (0.87) [12], in keeping with hypotheses that skin pigmentation variation has been subject to strong selection [13, 14].

A phenotypic trait of recent considerable interest is the level of gene expression (or RNA abundance), as it represents the initial link between genotype and other phenotypes, and hence is the logical place to begin evaluating the relative influence of genotype, environment and non-neutral evolution on phenotypic variation. Previous studies [15-21] have analyzed gene expression in lymphoblastoid cell lines from up to eight global populations derived from the International HapMap Project [22], and estimated that between 4.5% - 29% of genes are differentially expressed among groups. Four of these studies have estimated a between-population component of expression variation [17, 19-21]. Specifically, when considering CEPH European (CEU) and Yoruba from Ibadan, Nigeria (YRI), the first of these studies estimated that 15% of expression variation was observed among groups, suggesting that expression variation mirrors genetic variation and hence is largely neutral [17]. A subsequent study [20] found a similar median estimate of 12% for the among-group variation in expression. However, after accounting for non-genetic factors that estimate was reduced to 5%. Another attempt to reduce non-genetic factors influencing expression variation obtained a median estimate of 0.7% between CEU and YRI samples [19], while the most recent study estimated 3% of the expression variation is found among groups [21]. It may be crucial to correct for non-genetic factors for these specific samples as they were collected at various times in the past, transformed into cell lines, and maintained in culture for up to 20 years [15, 22, 23]. Yet given the range of estimates, the question remains, what proportion of total gene expression variation is found among groups, especially for native tissues rather than cell lines?

Here, we provide one of the first studies of among population gene expression variation in a natural tissue (namely, placentas) [24]. We chose placentas rather than more-easily obtained blood samples because gene expression in blood is influenced by the age of the individual [25] and the time of the day when samples are taken [26], whereas all placentas were obtained at the same “age” and “time” – namely, birth of the child. Additionally, the placenta is an important organ due to the fetal-maternal interplay and its critical role in fetal growth and development. Placentas were obtained from a single hospital during a six-week time period from four groups: African-Americans (AF), European-Americans (EU), South Asian Americans (SA), and East Asian Americans (EA). We emphasize that although we have tried to minimize environmental variation by sampling from a single hospital over a short time period, any differences in gene expression among these four groups will reflect both differences in genetic ancestry as well as systematic differences in their individual environments. However, we also incorporated biological and environmental factors into our model of gene expression to explicitly dissect the contributing variation that individual biology and environmental elements, such as diet, may have on expression variation.

A complication in the study of native tissues, such as placentas, is their cell type heterogeneity, and their spatio-temporal expression variability [27-29]. Thus, any one dissection of a complex tissue is but a single snapshot of the stochastic variation observed in expression abundance in that tissue space and in that moment in time. We therefore sampled each placenta twice to explicitly measure variation within a single placenta, to estimate the contribution of cell-type heterogeneity and spatial variability to inter-individual variation.

## Results

Of the 159 million high quality reads obtained, 117 million mapped to annotated exons. An average of 1.46 million exon-mapped reads were obtained for each library (sample replicate), corresponding to an average of 2.9 million exon-mapped reads for each individual (Figure S1 in Additional File 1). There was at least one mapped read for each library at 13,156 genes, including but not limited to 11301 protein-coding genes, 801 psuedogenes, 893 long noncoding RNAs (lncRNAs), and 40 small RNAs, which includes 21 pre-miRNAs. Expression levels were normalized (variance stabilized) using protocols described in the DESeq2 package [30]. Pearson’s correlation coefficient for each pair of sample replicates was 0.98 ± 0.005, yielding an r-squared value of 0.96 ± 0.01. Data quality was further evaluated by validating the expression profiles of three genes by rt-qPCR, a mean Pearson’s r of 0.74 ± 0.07 was observed between the expression values measured by RNA-sequencing vs. rt-qPCR (Figure S2 in Additional file 1). Thus, based on both sample replicates and an independent method of measuring expression abundance, the data we obtained accurately provide an accurate measurement of RNA transcript abundance.

### Total Gene Expression Structure

To determine if inter-individual gene expression variation was larger than intraindividual variation, and if individuals cluster by ancestry, a sample-by-sample correlation matrix was calculated and a hierarchical clustering dendrogram of all libraries was produced (Figure 1A). We observed that 74 of the 80 dissection replicates clustered together, consistent with the correlation results and indicating that intra-individual variation tends to be smaller than inter-individual variation. The three individuals whose dissection replicates did not pair were subsequently removed from all further analyses under the assumption that their lack of pairing was the product of dissection and/or processing error.

**Figure 1.**
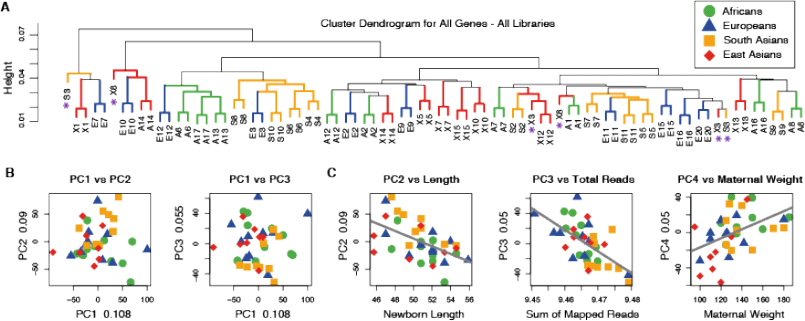
Overview of total gene expression variation at 13156 expressed genes. (A) A cluster dendrogram of libraries based on the following expression distance between each pair of libraries: 1-abs(r), where r is Pearson’s correlation coefficient for expression levels across all genes. Individual libraries and branches are colored to designate their group affiliation; asterisks indicate three pairs of replicate libraries that do not cluster together. (B) Scatter plots of the first three principle components (PC) using data from individuals and all genes. The explained proportion of variation is annotated on each axis. (C) Scatter plots between the first few PCs and correlated explanatory variables.

An additional observation from the sample correlation dendrogram is the lack of clustering of individuals with the same ancestry. To further evaluate this a principle component (PC) analysis reveals that in contrast to what is commonly observed for genetic data [31-33] there is no evident structure in this cellular phenotype that corresponds to groups (Figure 1B). However, when the PC loadings for each individual are tested for correlations with other aspects of the data (Figure 1C), PC2 is correlated with fetal length at birth (r = −0.54, Bonferroni p-value = 0.007), PC3 correlates with the sum of mapped reads (r = −0.62, Bonferroni p-value = 0.0005), and PC4 correlates with normal maternal weight (r = 0.46, Bonferroni p-value = 0.045). Additionally, analysis of genes that correlate with the loadings from the first 3 PCs [34, 35] reveals enrichment in hundreds of gene ontology categories, particularly molecular function (GO:0003674), biological process (GO:0008150), binding (GO:0005488) and their sub-categories (Additional file 2 - Tab A), as well as numerous KEGG pathways (Additional file 2 - Tab B) highlighted by the most enriched KEGG pathway, namely 01100:Metabolic Pathways (adjusted p-value = 2.9e-05). Overall, it appears that total transcriptome variation is largely influenced by factors other than group affiliation (e.g., population), and that transcript variation hence does not parallel expected patterns of genetic structure for these groups [32, 36].

### An Apportionment of Gene Expression Variation

Total variance in expression at each gene was apportioned among groups (Mst & Nst), among individuals within groups (Mit and Nit) and among dissection replicates (or within individuals, Met and Net). An analysis of variance (ANOVA) at each gene was performed to apportion the variation and two components of the data were used to derive the apportionment estimates - the additive components of variances and the sums of squares estimates (see the Methods section for details on these models.). Under this framework we are able to model all groups simultaneously as well as model populations in pairs. Assuming a model with four populations, the variance (Mst, Mit. Met) and variation parameters (Nst, Nit, Net) are highly correlated across genes (Mst:Nst, r = 0.97; Mit:Nit, r = 0.95; Met:Net, r = 0.99; p = 2.2e-16; Figure S3 in Additional File 1), even though their distributions and mean estimates are quite different (Figure 2A-C). The uniqueness of the variance parameters (M*t) reflects the specific manner in which these values are derived – that is, by the additive component of variance from the expected mean squares in this type I hierarchical ANOVA (see: Table S1 and S2 in Additional file 1). Given the correlation among parameter estimates and the lack of zero values in the sums of squares approach (Figure S3 in Additional File 1), we focus on the variation or variability parameters Nst, Nit and Net. On average we find that 33.2% of the variability in gene expression is found among populations of cells within a single tissue (Net, permutation of reads among replicates, p = 0.22), 58.9% of the variability is among individuals within groups (Nit, permutation of libraries among individuals within groups, p = 0.048) and 7.9% of the variability is among groups (Nst, permutation of individuals among groups, p = 0.24) (Figure 2 B-C). These estimates indicate that even though inter-individual variation is, on average, the largest component of expression variation, intra-individual variation cannot be ignored in measuring cellular phenotypes. Similarly, while among group expression variation does not, on average, reach the levels of structure seen at the genetic level, the group component does detectably influence expression variation, particularly at a subset of genes, which we explore below.

**Figure 2.**
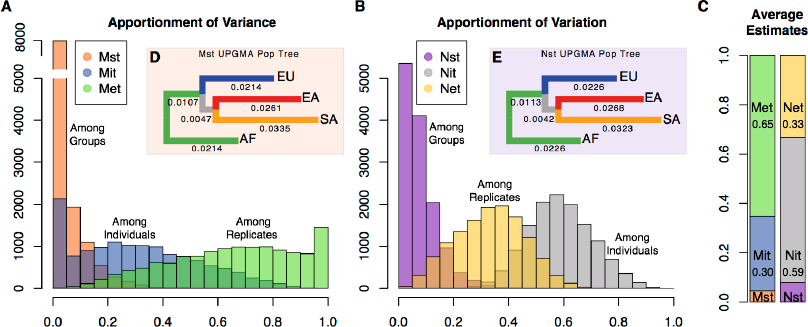
Apportionment Summaries. (A) The distribution of variance apportionments derived from the additive component of variance estimates. (B) The distribution of variation apportionments derived from the sum of squares. (C) Mean estimates for each apportionment parameter using both the variance and variation. (D) A dendrogram of weighted mean population distances derived from the Mst parameter. (E) A dendrogram of weighted mean population distances derived from the Nst parameter.

When modeling expression variation in a pairwise manner, mean estimates are similar to those observed in the four-population analysis (Table 1). However, among group variation (Nst) ranges from 0.045 (for AF:EU) to 0.062 (for EA:SA). A dendrogram was constructed using mean pairwise Nst distances (Figure 2D-E). We find that the data are congruent with expectations from genetic data [36], with the exception that SA tend to be the most distant group.

**Table 1.** Apportionment estimates. Variation apportionment estimates for each pairwise population comparison and the single model that evaluates all populations at once (4pop). Population annotations are as follows: AF = African American, EU = European Americans, SA = South Asian Americans, EA = East Asian Americans. Nst = porportion of total variation explained by groups, Nit = proportion of total variation explained by individuals within groups, Net = proportion of total variation explained by dissection replicates within individual and by error, Nis = proportion of inter-and intra- individual variation explained by variation observed among individuals.

|  | **Nst** | **Nit** | **Net** | **Nis** |
| --- | --- | --- | --- | --- |
| **4pop** | 0.079 | 0.589 | 0.332 | 0.641 |
| **AF:EU** | 0.045 | 0.629 | 0.326 | 0.66 |
| **AF:SA** | 0.061 | 0.607 | 0.331 | 0.649 |
| **AF:EA** | 0.059 | 0.599 | 0.343 | 0.638 |
| **EU:SA** | 0.054 | 0.617 | 0.329 | 0.653 |
| **EU:EA** | 0.049 | 0.611 | 0.34 | 0.643 |
| **SA:EA** | 0.062 | 0.589 | 0.348 | 0.63 |

### Mean expression and apportionment estimates

The mean expression of each gene is significantly correlated with the residual (or intra-individual) sum of squares estimate (Pearson’s r = 0.60, p ≪ 0.001). This illustrates that as mean expression increases, variation in mRNA abundance among our sample replicates also increases. As such, we estimate that mean expression explains 36% of the variation in our error sum of squares. However, the among group (r = 0.018, p = 0.034) and among individuals within groups (r = −0.029, p = 0.001) sums of squares are more weakly correlated with mean expression. Consequently, the apportionment parameters are correlated with mean expression with coefficients of −0.446, 0.388 and 0.166 for Net, Nit and Nst (p ≪ 0.001), respectively. The proportion of variation explained by mean expression for each apportionment parameter is thus 20%, 15% and 2.7% for Net, Nit and Nst, respectively. This suggests that mean expression is having a modest influence on parameter estimates, and the acquisition of more reads will not greatly influence the apportionment estimates.

### Differential Gene Expression Among Individuals

The proportion of genes that vary significantly among individuals in expression levels was analyzed via a F-ratio test between inter-individual and intra-individual variance. We observed that 5880 genes, or 44.5% of all genes (at an FDR 5%), exhibited significant among individual, within group variation. Additionally, fitting two linear models to the data (a null model and a second model that includes individuals as an explanatory variable), followed by a Chi-squared test of model fitting, results in 5491 genes (41.7% of all genes) with significant inter-individual variance (at an FDR 5%). There is an 84% overlap between the significant genes in both analyses. We estimated the proportion of within-group variation explained by inter-individual variation with the parameter Nis (SSb/SSb+SSe; see Methods). On average 64% of the within-group variation is attributed to individuals, indicating substantial inter-individual variation. Those genes that are significantly differentially expressed (DE) among individuals, as determined by the F-ratio test, have a minimum Nis value of 0.65. To determine if there may be significant variation attributed to intra-individual variation at some loci, we inverted the F-ratio test by placing the intra-individual mean squares in the numerator and inter-individual mean squares in the denominator, but observed no significant loci after Benjamini-Hochberg correction. Overall, this illustrates that there is substantial interindividual variation in gene expression variation.

### Differential Gene Expression Among Groups

Three different methods were used to identify and quantify genes that may be differentially expressed among human groups: two published methods (DESeq [30] and tweeDESeq [37]), and a permutation of the hierarchical ANOVA. The two published methods can only compare two groups at a time, while permutations of the hierarchical ANOVA permit the analysis of two or more groups simultaneously.

While there is marked variation in the number of DE genes that each method identified, there are consistent trends (Table 2). For example, the relative proportion of DE genes for each pair of populations were correlated between methods (Pearson’s r = 0.927, p < 0.008) and comparisons that included South Asians tended to have the most DE genes for any one group. Further, 99% and 92% of the genes identified as DE by the DESeq and tweeDESeq methods respectively were also identified as DE by the permutation method. In the permutation analysis, the cutoff Nst value for DE genes differs slightly depending on the groups being compared but averages out to an Nst estimate of at least 0.326. The reduced number of DE genes identified with the DESeq and tweeDESeq methods is because both methods are model-based analyses with specific tests and false discovery correction of differential expression. The permutation method presented here simply identifies extremes in the observed data that are difficult to explain by random chance.

**Table 2.**
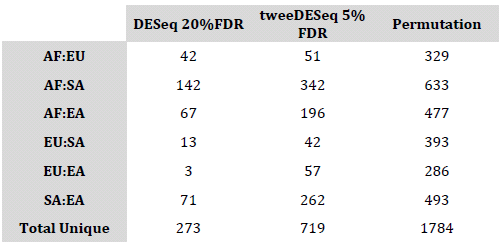
Number of Differentially Expressed Genes. The estimated number of differentially expressed genes between each pair of populations, determined using three different methodologies.

|  | **DESeq 20%FDR** | **tweeDESeq 5%**<br>**FDR** | **Permutation** |
| --- | --- | --- | --- |
| **AF:EU** | 42 | 51 | 329 |
| **AF:SA** | 142 | 342 | 633 |
| **AF:EA** | 67 | 196 | 477 |
| **EU:SA** | 13 | 42 | 393 |
| **EU:EA** | 3 | 57 | 286 |
| **SA:EA** | 71 | 262 | 493 |
| **Total Unique** | 273 | 719 | 1784 |

To determine the potential biological relevance of the genes identified as DE, we tested for enrichment in GO and KEGG pathways. When testing the union of all pairwise permutation DE genes (1784 DE genes), we observed enrichment in 15 KEGG pathways and 371 GO categories at a moderate-confidence FDR of 20% (5 KEGG and 201 GO at a high-confidence FDR of 5%) (Table 3, Additional file 3 - Tab A). In general, KEGG and GO enrichments indicate that genes involved in cellular signaling, immune response, tissue and organ development, and metabolism pathways are DE among groups.

**Table 3.** Enriched pathways for the pairwise union of all DE Genes. Shown are the KEGG pathways and GO categories observed to be enriched when using the union of genes identified as DE (1784 genes) in each pairwise comparison found in Table 2 “Permutation”. The table provides the category identifiers (in the case of GO the ontology: CC, cellular component; BP, biological process; MF, molecular function), the associated term or brief description, and the Benjamini-Hochberg adjusted p-value.

| category | KEGG TERM | padj | GO Category | Ontology | GO Term | padj |
| --- | --- | --- | --- | --- | --- | --- |
| 4514 | Cell adhesion molecules (CAMs) | 0.0046 | GO:0032501 | BP | multicellular organismal process | 6.64E-10 |
| 5144 | Malaria | 0.0059 | GO:0044707 | BP | single-multicellular organism process | 4.57E-09 |
| 4142 | Lysosome | 0.0059 | GO:0006950 | BP | response to stress | 5.06E-09 |
| 5143 | African trypanosomiasis | 0.0143 | GO:0044699 | BP | single-organism process | 1.27E-08 |
| 4512 | ECM-receptor interaction | 0.0238 | GO:0044763 | BP | single-organism cellular process | 9.29E-08 |
| 4145 | Phagosome | 0.0678 | GO:0050896 | BP | response to stimulus | 1.01E-07 |
| 380 | Tryptophan metabolism | 0.0751 | GO:0004872 | MF | receptor activity | 2.07E-06 |
| 5020 | Prion diseases | 0.0751 | GO:0002376 | BP | immune system process | 2.07E-06 |
| 4610 | Complement and coagulation cascades | 0.0751 | GO:0007275 | BP | multicellular organismal development | 2.07E-06 |
| 5416 | Viral myocarditis | 0.1024 | GO:0032502 | BP | developmental process | 2.34E-06 |
| 4640 | Hematopoietic cell lineage | 0.1366 | GO:0007154 | BP | cell communication | 5.70E-06 |
| 5414 | Dilated cardiomyopathy | 0.1513 | GO:0023052 | BP | signaling | 6.39E-06 |
| 5150 | Staphylococcus aureus infection | 0.1513 | GO:0044700 | BP | single organism signaling | 6.39E-06 |
| 590 | Arachidonic acid metabolism | 0.1513 | GO:0048731 | BP | system development | 7.32E-06 |
| 480 | Glutathione metabolism | 0.1683 | GO:0006955 | BP | immune response | 7.32E-06 |

### Non-Neutral Gene Expression Profiles

Although it is difficult to determine if expression at a particular gene is evolving according to neutrality or under selection, we are able to identify expression profiles that conform to four specific patterns of selection: directional, balancing, stabilizing and diversifying. Importantly, these analyses do not test for deviations from neutrality, but rather identify genes that exhibit expression profiles consistent with selection on quantitative traits [38, 39]. Traits under directional selection are expected to exhibit shifts in mean expression among groups exemplified by greater among group variation relative to within group variation, and would hence be consistent with previously identified DE genes. Balancing selection is exemplified by high diversity or variation among individuals within a population but low variation among populations. Stabilizing selection results in low levels of expression variance among individuals while diversifying selection is reflected in high levels of expression variance among individuals. We identified genes that typify each selection profile using apportionment of variation estimates, estimates of total expression variance, and a series of permutations, as described in Materials and Methods.

Using data from the model fitting all four groups simultaneously, we observe that the among groups variation (log(SS_a_)) correlates positively with the among individuals variation (log(SS_b_), Pearson’s r = 0.579, p < 2.2e-16), in agreement with expectations under neutrality [40]. Additionally, the variation within individuals (log(SS_e_) also correlates positively with the among individuals variation (Pearson’s r = 0.46, p < 2.2e-16) and the among groups variation (Pearson’s r = 0.25, p < 2.2e-16) (Figure 3A). To estimate the proportion of the human placental transcriptome that may be consistent with neutral vs. non-neutral expectations, we performed a series of permutations (see Materials and Methods). We estimate that 64.8% of all genes are consistent with a neutral-drift model for a quantitative trait [38]. The most prevalent non-neutral profile of gene expression variation is stabilizing selection, which influences an estimated 26% of all genes, followed by directional (646 genes, 4.9%), diversifying (635 genes, 4.8%), and balancing (173 genes, 1.3%) selection (Figure 3B; see Additional file 5 for a list of all genes).

**Figure 3.**
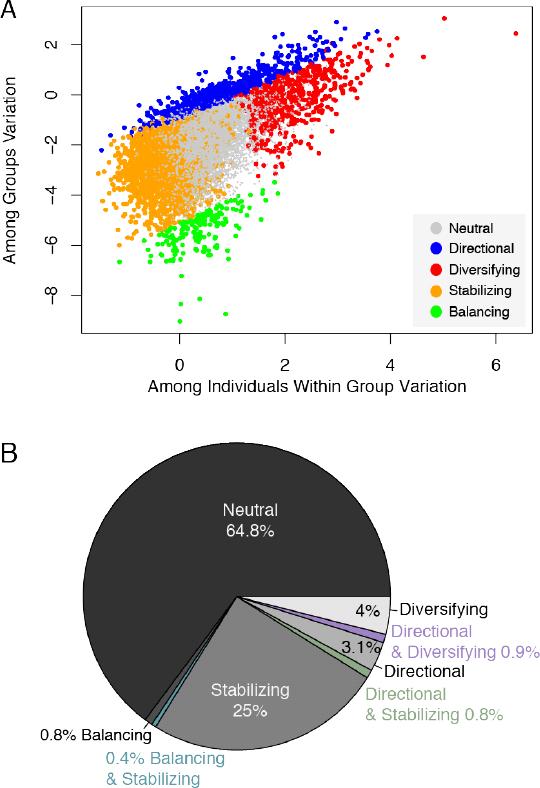
Evaluating neutral vs. non-neutral evolution of the human placental transcriptome. (A) A scatter plot of among group and among individual variation as measured by the log of the corresponding sum of squares. Genes that were identified as having patterns of variation consistent with neutrality or with directional, diversifying, stabilizing or balancing selection are color-coded. (B) A pie chart illustrating the proportion of genes consistent with a particular mode of evolution.

When each of these modes of selection are mapped onto the distribution of within-group and among-group variation (Figure 3A) we can identify near discrete sections of the distribution that reflect these observations. Interestingly, there are areas of the distribution where these modes of selection overlap (Figure 3B). For example, there is a small set of genes for which expression variation is both large among individuals (diversifying) and among groups (directional) (Figure 4A-B). Conversely, some genes have more constraint in total variance, consistent with stabilizing selection, and yet also have significant shifts in mean expression among groups, consistent with directional selection (Figure 4A and C). And finally, constrained inter-individual expression (stabilizing selection) can also occur with reduced among group variation (balancing selection) (Figure 4D).

**Figure 4.**
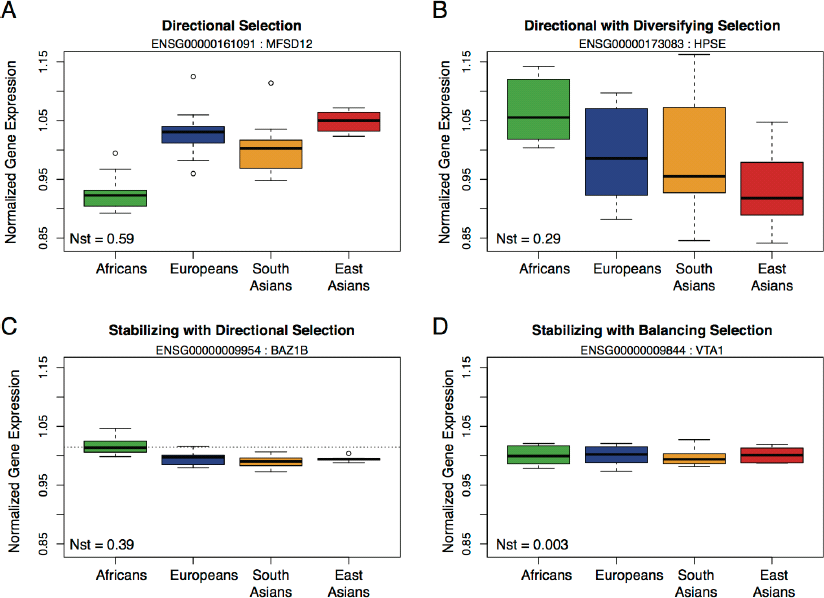
Boxplots of non-neutral expression variation. The y-axis of all plots illustrates the same range of expression. Each population is color-coded and the estimated Nst value for each gene is in the bottom left corner of each plot. (A) A gene consistent with directional selection. (B) A gene consistent with both directional and diversifying selection. (C) A gene consistent with both stabilizing and directional selection, with a dotted grey horizontal line to help view the shift in mean expression, while also presenting constrained among group, within individual variation. (D) A gene consistent with both stabilizing and balancing selection.

To determine if genes differentially expressed among groups, i.e. those with a pattern consistent with directional selection, could effectively recapitulate group ancestry, we used expression variation across all 646 directional genes (those identified when modeling all four populations at once) to generate a UPGMA tree and perform a principle component analysis. We observe that individuals form monophyletic clades consistent with population ancestry (Figure 5A). Additionally, increased levels of population structure were observed in the principle component analysis but are only fully discernable when viewing the first 3 PCs together (Figure 5B). PC1 tends to distinguish individuals of African ancestry from those of non-African ancestry, while PC2 tends to distinguish SA from EA and PC3 distinguishes Europeans from non-Europeans (Figure S4 in Additional file 1).

**Figure 5.**
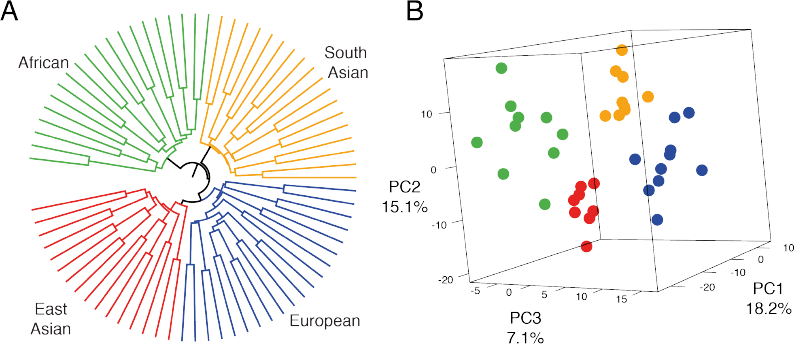
Population structure revealed by genes consistent with directional selection. (A) A UPGMA tree of expression distances among all libraries and individuals at genes consistent with directional selection. (B) A 3D scatter plot of the first 3 PCs based on variation in the 646 genes consistent with directional selection. The proportion of explained variation is annotated on each axis and each individual’s group affiliation is color-coded to match the annotation in plot A.

### Expression Variance, Genetic Diversity and Network Connectivity

The prevalence of genes that deviate from neutral-drift expectations, particularly those consistent with stabilizing selection, prompted us to hypothesize that inter-individual variance in gene expression must have a genetic component. Specifically, we hypothesized that genes with greater expression constraint would have greater genetic constraint. Additionally, genes exhibiting large inter-individual expression variances may allow, through relaxed constraint or by necessity, a relative excess of variation. To evaluate this hypothesis, we tested for a correlation between expression variance and pairwise genetic diversity. Pairwise genetic diversity (π) was calculated for each gene, controlling for gene length [41], for 3 populations from the 1000 Genomes data: CEU-Northern Europeans, ASW- African Americans from the S.W. USA, and CHS – Han Chinese from Southern China. We chose these three populations as they are the best available proxies for our sampled individuals. When diversity is compared from each population to expression variance, we observe a significant positive correlation (ASW: r = 0.213; CEU: r = 0.189; CHS: r = 0.177, p≈0, Figure S5 in Additional file 1). In addition, expression variance also correlates with Tajima’s D values (ASW: r = 0.179; CEU: r = 0.129; CHS: r = 0.132, p≈0). These observations indicate that total expression variance has a small (r-squared ≈ 0.04) albeit significant genetic and thus heritable component. Another factor that may influence expression variance is the number of interacting partners a gene has. Previous work on gene-networks has illustrated that the degree of connectivity (number of interactions) influences the rate of molecular evolution [42]. Here, using data from BioGrid we tested if the number of interacting genes also influences the expression variance of a gene (Figure S6 in Additional file 1). Indeed, we observe a weak tendency for the expression variance to increase as the number of interacting genes decreases (Pearson’s r = −0.28, p≈0).

To evaluate how both genetic diversity and connectivity may together influence gene expression variance we built an ANOVA model setting the coefficient of variation in gene expression as the response variable, and setting gene diversity and connectivity as explanatory variables with interaction. Each component of the model significantly influenced expression variance (diversity p≈0; connectivity p≈0; interaction p = 0.029) explaining an estimated 4.3%, 2.3% and 0.07% of the total variance in expression variance.

### Gene Co-Expresssion Modules & Functionality of Selection Categories

To determine if the sets of genes corresponding to the four non-neutral modes of evolution have a coherent biological effect, we tested for evidence of co-expression networks and enrichment in GO gene ontology terms and KEGG functional pathways. No enrichment was observed for genes consistent with a pattern of balancing selection. The results from the three other non-neutral modes are presented below.

Overall, genes consistent with directional selection (646 genes) were enriched in 145 GO categories and 6 KEGG pathways at an FDR of 20% (70 and 0 respectively at an FDR of 5%). They are associated with extracellular and membrane regions, response to stress, infectious disease, signaling, binding and metabolism pathways and categories (Additional file 3 – Tab B). Six coexpression modules were identified that form compact coexpression networks, but also interact with each other through a reduced number of loci (Figure 6A,B). The only individual module that is enriched for a particular set of functions is module 6 (red Module in Figure 6A). This is the smallest module, containing just 54 genes, but at an FDR of 20% this module is enriched for 110 GO categories (52 at FDR 5%, Additional file 3 – Tab C), and 15 KEGG pathways (7 at FDR 5%, Additional file 3 – Tab D). These genes are principally involved in defense and immune response but are also associated with vitamin absorption and digestion, and arachidonic acid metabolism, a key fatty acid.

**Figure 6.**
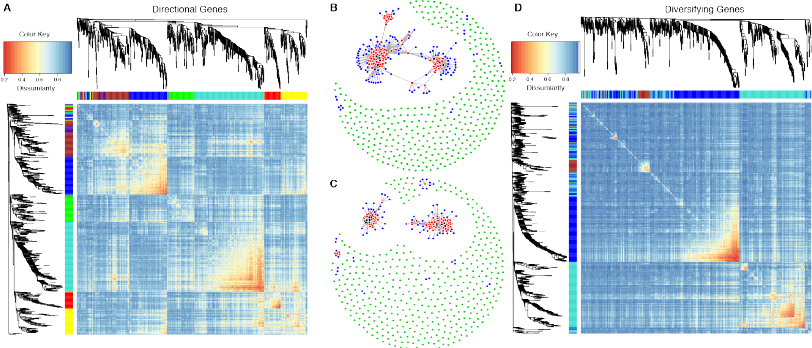
Co-expression heatmaps and networks. Heatmaps of gene × gene expression correlations for genes under directional selection (A) and diversifying selection (D), respectively. Each row and column is the same set of genes, annotated by the same cluster dendrogram of gene expression distance. Additionally, each row and column is color-coded to its associated gene co-expression module. In the heatmap plot itself, the color red indicates more similar co-expression and blue indicates greater dissimilarity. Gene co-expression networks for genes under directional selection (B) and diversifying selection (C) are also presented. Nodes of interaction were only created for genes which present significant co-expression at an FDR of 1%. Black nodes are genes with at least 32 significant interactions. Red nodes are genes with at least 7 significant interactions. Blue nodes are genes with at least 2 significant interactions. Green dots are genes with no significant interactions at an FDR of 1%.

To evaluate if the enrichment observed here is the product of unique expression in a particular population or variation across all groups, we partitioned all directional genes by their expression profiles using k-means clustering. When partitioning the expression profile data into two groups (k=2), we observe two opposing profiles where expression is lowest in Africans, highest in South and East Asians, and intermediate in Europeans (cluster 1) or highest in Africans, lowest in South and East Asians, and intermediate in Europeans (cluster 2) (Figure 7, row K2). Enrichment tests for these two clusters reveal that only cluster 1 exhibits any enrichment, with ontology and pathway enrichment consistent with those observed above. This observation would be consistent with a hypothesis of adaptive responses in non-African populations during migrations out of Africa. However, when the data are partitioned into more clusters (k=6), there is no ontology or pathway enrichment for those clusters that accentuate the expression differences between Africans and non-Africans (Figure 7, row K6, clusters 4 and 5). Note that we chose a K of 6 for this particular analysis because it is the first K that uniquely separates African from non-African populations in both an up-regulated (cluster 4) and down-regulated (cluster 5) manner. Results for K2 through K8 can be found in Additional file 1, Figure S7. Interestingly, it is rather cluster 1 (Figure 7, row K6), with elevated expression in South Asians relative to the other groups, that harbors the entire enrichment signal. These 111 genes are enriched at an FDR of 20% in 19 KEGG pathways (8 at FDR 5%) and 320 GO categories (136 at FDR 5%). Again, they are mostly involved in immune response and metabolism, consistent with the observations above (Additional file 3 – Tab E).

With diversifying genes, 3 coexpression modules (Figure 6D) were identified and 2 highly integrated networks along with 2 smaller networks (Figure 6C), consistent with the coexpression modules, were observed. Each module was enriched in numerous GO ontology terms (Additional file 3 – Tab F) and KEGG pathways (Additional file 3 – Tab G) with both unique and overlapping functions. Module 1 (Figure 6D, cyan) is enriched in 546 GO ontology terms and 22 KEGG pathways at an FDR of 20% (222 GO and 8 KEGG at FDR 5%) and involved in numerous areas of biology including growth, development, signaling, metabolism and disease. Module 2 (Figure 6D, blue) is enriched in 131 GO ontology terms and 3 KEGG pathways at an FDR of 20% (35 GO and 2 KEGG at FDR 5%) and involved with binding and receptor interaction, specifically cytokine-cytokine receptor interaction and neuroactive ligand-receptor interaction. Module 3 (Figure 6D, dark red) is enriched in 378 GO ontology terms and 12 KEGG pathways at an FDR of 20% (132 GO and 9 KEGG at FDR 5%) and associated with disease and signaling pathways. The union of all diversifying genes reveals ontological and functional enrichment consistent with the above data (Table 4, Additional file 3 – Tab H).

**Table 4.**
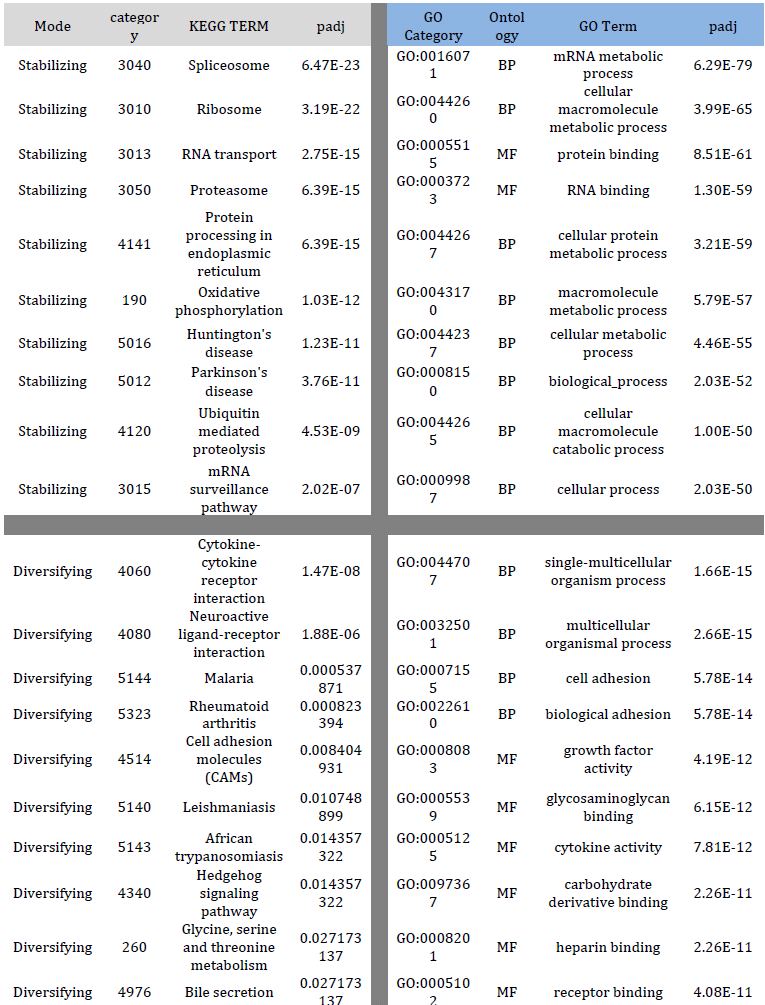
Enriched pathways for stabilizing and diversifying genes. The table provides the selective mode, the category identifier (as in Table 3), the associated term or brief description, and the Benjamini-Hochberg adjusted p-value.

**Figure 7.**
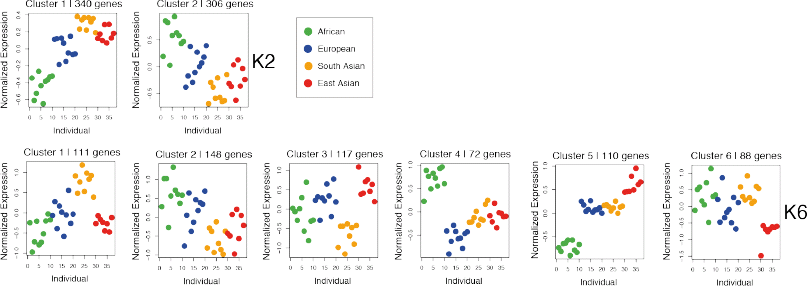
Expression levels for genes consistent with directional selection. Each dot represents an individual spaced across the x-axis and with mean normalized gene expression on the y-axis. The results of the cluster analysis are illustrated for two clusters (K2) and for six clusters (K6). Individuals are color-coded with respect to their associated group.

Stabilizing genes formed 4 coexpression modules that, as a unit (Additional file 3 – Tab I), are associated with 1245 GO ontology terms and 51 KEGG pathways at an FDR of 20% (898 GO and 39 KEGG at an FDR of 5%) and are involved with basic, largely intracellular, processes (Table 4). These include association with the splicesome, ribosomes, RNA transport and protein processing. But they are also associated with neurological diseases such as Huntington’s, Parkinson’s and Alzheimer’s disease. Finally, there are also associations with bacterial infection, hepatitis C, T-cell signaling and cancer pathways. Individually, each module has a unique functional composition, but there is overlap at varying degrees for a few key pathways that include basic intracellular functions and associations with neurological diseases (Additional file 3 – Tab J and K).

### The influence of biological traits on gene expression

Along with population ancestry, several anthropometric and dietary traits were also collected from each individual, to evaluate their association with expression variation. Starting with the model of gene expression used previously, which included technical (number of mapped reads and RNA quality) and population factors (group and individual), eight additional traits were added: sex of the child, weight of the child, length of the child, birthing manner (cesarean or vaginal), maternal age, maternal body mass index, whether or not the mother drinks alcohol (outside of the pregnancy), and whether or not the mother is a vegetarian (see Materials and Methods for model details). Note that each new trait being modeled is a measure of inter-individual variation. The significance for each factor was determined by an F-test (FDR of 5%) using the mean square estimates of each factor over the residual (intra-individual variation).

On average each factor explained roughly 2% of the variation in the data, with intra-individual (32%) and inter-individual (41%) variation accounting for most of the variance; among group variation explained 6.3% (Figure 8). As expected the vast majority of variation explained by each of the new explanatory variable was previously explained by variation among individuals, thus the reduction in the Nit estimate from 0.59 (Nit, Figure 2C), to 0.41 (Figure 8). All factors were enriched in no less than 59 GO ontology (Additional file 4 – Tab A) terms at an FDR of 5% and all but 3 factors (RIN, sex and length) were enriched in at least one KEGG pathway at an FDR of 5% (Additional file 4 – Tab B). Importantly, the significance for all factors was dependent on the within group-among individual variation (Nit) and the mean expression of genes (Figure S7 in Additional file 1). As such, if a gene previously exhibited no significant variation among individuals in our simple model of gene expression then it did not exhibit any significant variation among any of the eight additional factors in our full model. Thus, all of the GO ontology terms and KEGG pathways observed for each of the new factors are simply a subset of those previously associated with variation among individuals, which was enriched in 104 KEGG pathways and 2720 GO ontology terms at an FDR of 20% (65 KEGG, 1729 GO at an FDR of 5%). On the technical side, genes that correlated with the number of mapped reads were overwhelmingly those that are highly expressed and associated with pathways such as Ribosome (KEGG 03010; adjusted p = 4.75e-23). Such technical artifacts are known to be an issue with this technology and are precisely why the number of mapped reads and RNA quality (RIN) values were included as leading explanatory variables in all models of gene expressions [43]. See Additional file 4, for all GO and KEGG enrichment data for each trait.

One striking observation from the trait model fitting was that newborn weight was associated with three cancer pathways and the hematopoietic cell lineage pathway. This observation is consistent with reports of newborn birth weight being associated with increased risks of childhood leukemia [44, 45]. Are the genes associated with this effect being down regulated as birthweight increases, or are they being up regulated? To evaluate this specific example and all other associated trait enrichments we partitioned the correlations between gene expression and the trait by the direction of their effect and then re-evaluated pathway associations (Figure 9, Additional file 4 – Tab C). The results indicate large coordinated changes in expression for each factor. For example, as newborn birth weight increases there is a decrease of expression in genes associated with the hematopoietic cell lineage, cancer pathways, bile secretion, dilated cardiomyopathy and vascular smooth muscle contraction, but genes associated with protein processing in the endoplasmic reticulum increases. Further, individuals who normally consume alcohol have decreased expression in pathways such as glycolysis and fat digestion. Placentas from female children have increased expression in protein digestion, ECM-receptor interaction, amoebiasis, and focal adhesion. Placentas from Cesarean births exhibit decreased expression in glycolysis, protein processing in the endoplasmic reticulum and antigen processing. As a final example - as maternal body mass index increases there is a correlated increase in expression for genes involved in staphylococcus aureus infection, complement and coagulation cascades and systemic lupus erythematosus pathways. These data, as presented in Figure 9, illustrate the correlated effect that gene expression changes may have on specific functional pathways and by inference on the physiology of an organ or individual.

**Figure 8.**
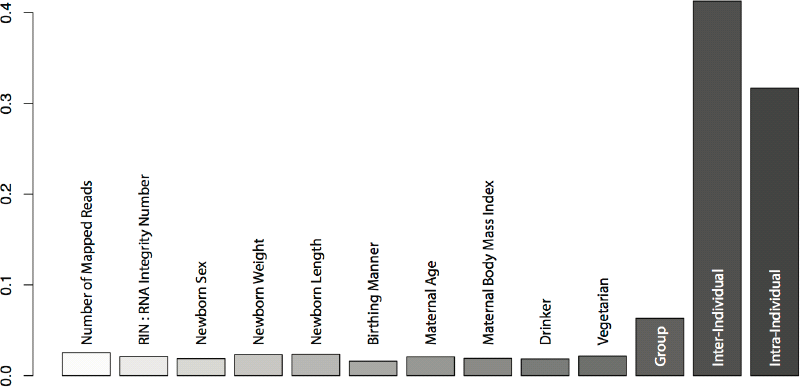
Apportionment bar plot. Each gene was fit to a single model accounting for 13 explanatory variables and the proportion of variation explained by each variable was estimated using the sum of squares approach.

**Figure 9.**
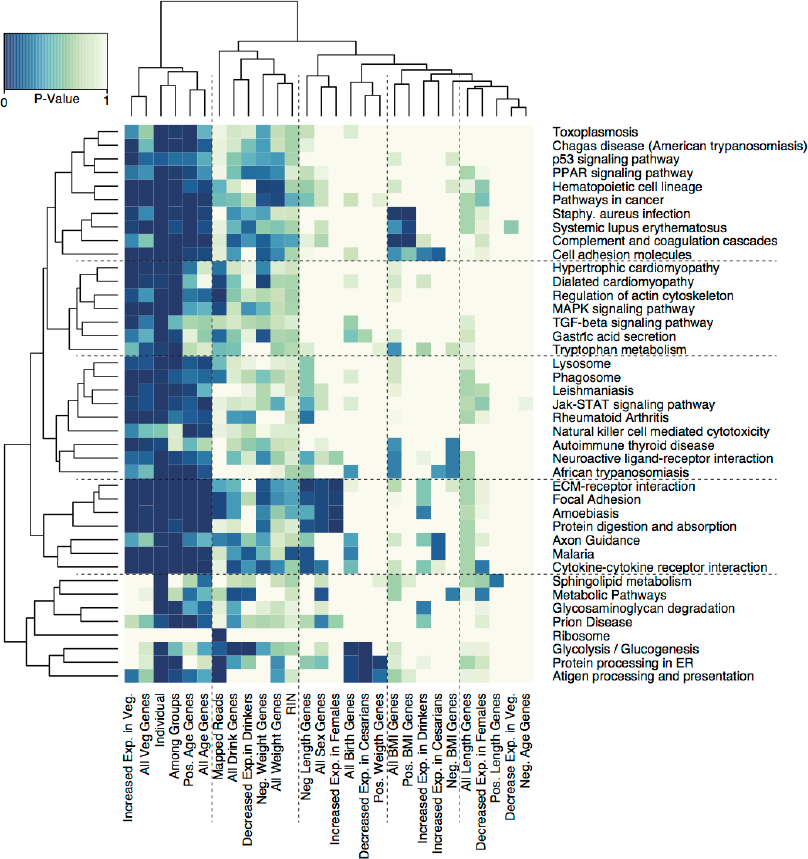
Enrichment heatmap. A heatmap of Benjamini-Hochberg adjusted p-values for the association between each explanatory variable (x-axis) and KEGG pathway categories (y-axis). To be included in the heatmap a KEGG pathway had to be associated with at least one explanatory variable at an FDR of 1%. Additionally, each explanatory variable was partitioned by the direction of its association with gene expression. For example, the variable “All Veg. Genes” annotates all genes that demonstrated a significant vegetarian diet effect, while the variable “Increased Exp. in Veg.” annotates those vegetarian diet associated genes whose expression profile increased relative to non-vegetarians. Similarly “Pos. Age Genes” annotates all genes that significantly correlated with maternal age in a positive manner.

## Discussion and Conclusion

Using a population genetics framework, we have demonstrated that both intra- and inter-individual variation account for the vast majority of total gene expression variation. Significantly, intra-individual variation in gene expression cannot be ignored in evaluating expression variation, consistent with studies of single cell gene expression that have illustrated the stochastic nature of expression variation [28]. While this is particularly true for the placenta it also holds true for other tissues [46, 47]. If intra-individual variation is not measured it will be erroneously attributed to inter-individual variation, thereby inflating estimates of interindividual variation.

Gene expression profiles were dissected to evaluate the impact that non-neutral evolutionary forces may play in shaping expression variation. We observed that the majority of placental expression variation is consistent with a neutral-drift model, but an estimated 35% of the placental transcriptome is influenced by selection. Stabilizing selection plays a large role on transcriptome variation maintaining significant regulatory control over some 25% of the genes. Genes influenced by stabilizing selection are largely limited to basic intra-cellular functions. In contrast, the ∼4% of genes influenced by diversifying selection typically encode extra-cellular proteins involved in cell signaling, metabolism and immune pathways. Interestingly, the genes influenced by directional selection span the range of inter-individual variation observed, overlapping with profiles consistent with stabilizing and diversifying selection (Figure 3A). That is, directional selection can act on any gene regardless of the range of inter-individual variance. Therefore, measurements of fold change in gene expression that do not account for total expression variance can be misleading (Figure 4). Additionally, we find that expression diversity correlates with genetic diversity, substantiating a role for genetic selection in influencing interindividual expression variation.

Among group variation in placental gene expression averages out to an Mst of 0.045 (Nst = 0.079), which is less than that found for human genetic variation (Fst = 0.111) [36]. This suggests that placental transcriptome variation among groups is more similar than genetic variation alone would predict. Our estimates and thus our conclusions are certainly influenced by the accuracy with which we can measure and apportion variation for such a dynamic and quantitative trait. However, these estimates are qualitatively consistent with similar recent estimates of among group variation derived from lymphoblastoid cell lines [19, 21].

Interestingly, where significant variation in expression level is manifested among human groups, it associates with genes pivotal to placental biology, fetal growth and fetal development. This includes cell-cell interaction pathways like cell adhesion molecules [48], arachidonic acid metabolism [49], tryptophan metabolism [50], and immune response pathways including malaria, which is known to present serious health risks to the fetus [51, 52]. While these inter-individual and inter-group observations are of potential interest for clinicians and biologists, a crucial point concerning the evolutionary consequences of these observations is heritability. In both the pairwise population analyses and k-means clustering of directional genes, individuals of South Asian ancestry appear to have experienced both the greatest change and the most biologically specific changes in placental gene expression. However, the mothers of all of these individuals were born in South Asia, and it is unknown how long they resided in the sample location prior to sampling. Thus, whether these observations are the product of a heritable, evolutionary adaptive response or the product of these particular individuals being exposed to an individually novel environment and presenting a plastic response cannot be determined. Nonetheless, those genes with expression profiles consistent with models of directional selection exemplify genes and pathways that may be most frequently targeted during adaptive responses to novel environments.

We have also demonstrated that by incorporating biological trait variation in models of gene expression, we can identify genes and pathways that have correlated changes with the modeled traits. By evaluating the direction of the correlated change and combining this information with biological and/or clinical information, this framework allows the potential influence of the trait to be dissected. For example, pre-pregnancy alcohol consumption is associated with the regulation of essential pathways like glycolysis/glycogenesis.

Finally, these observations provide a first insight into human, in vivo, gene expression variation among populations of cells within a tissue, among individuals and among continental groups. The model of gene expression variation presented here is adaptable to any system and the apportionment parameters based on the sums of squares provide a set of stable statistics that can be compared across studies. Importantly, classifying genes into selection categories is difficult as there are a number of assumptions involved. We stress that no formal tests of selection were performed in this study. Instead, we presented a framework to identify genes with expression profiles that are consistent with theoretical expectations of selection on a quantitative trait. Hopefully, this work will provide a foundation for the development of a neutral theory of gene expression in which formal tests of selection may be conducted. Further, we note that precision in the apportionments can be strengthened by increasing the number of sequencing reads, adding technical replicates, and increasing the number of both tissue replicates and individuals. Notably, only a single complex tissue was evaluated in this study - additional biological and evolutionary insight can be gained by studying other tissues or, as single-cell transcriptomic methodologies become more mature, specific cell types. In addition, sampling individuals of similar ancestry at multiple locations would allow one to estimate the influence of both environment and ancestry on expression variation. The framework and methodologies present here provide a foundation for further such studies of transcriptome variation.

## Methods

### Ethics Statement

All placentas were collected in October - November 2006 at Northside Hospital in Atlanta, Georgia, with the approval of the Northside Hospital Institutional Review Board (NSH #804) and with the written informed consent of the donors.

### Samples

A total of 66 human placentas were collected and processed within an hour of delivery from both natural and cesarean births. Placentas were quartered, wrapped in aluminum envelopes, and immediately snap frozen in liquid nitrogen. All samples were stored at −80°C prior to shipping on dry ice to the Max Plank Institute in Leipzig, Germany, where they were again stored at −80°C. Each contributing family completed a questionnaire which asked for self-described ancestry and birthplace going back three generations and anthropometric, health and life-style questions about the mother including: height, weight, weight at full term, number of pregnancies, number of children, smoking status, alcohol intake, illness during pregnancy, chronic illnesses, medication taken during pregnancy, diet and any other volunteered information. Finally, the sex, weight, length and the delivery manner of the child were recorded. From this collection and the provided data, we selected 40 samples to include in the study. Samples were chosen only from those families with self-described ancestry from a single group, with no major illnesses during birth, and with the most complete questionnaires. The final 40 samples include ten samples each of African-American (AF), European-American (EU), South Asian-American (India; SA) and East Asian-American ancestry (Korea, China, Vietnam and Taiwan; EA). All SA individuals are first-generation immigrants, and all but one of the EA individuals are first-generation immigrants; the exception is a second-generation American.

### Dissections

Given the mosaic composition of the placenta, possible maternal blood/tissue contributions to any dissection, and previous observations that placental sample location influences expression variation [27], we produced tissue sample replicates for each individual. Tissue sample or dissection replicates were generated to quantify expression variation introduced in the dissection process. Specifically, tissue replicates quantify intra-individual variation in the form of (a) cell-type heterogeneity, (b) biological variation across a tissue and (c) temporal and stochastic variation in gene expression, thus allowing for a more accurate estimation of the variation found among individuals. From three of the four quarters of each placenta we dissected 100mg of centrally located villus parenchyma tissue (taking care to avoid decidua, chorion, or amnion tissue) from five nonadjacent locations, totaling 600 dissections. The five dissections from each quarter were pooled, resulting in three sample replicates from each placenta. Five non-adjacent dissections were taken in an effort to homogenize the cell-type composition among samples. All dissections were carried out on a sterilized steel plate situated on top of dry ice, thereby keeping the samples frozen at all times. Samples for dissection were chosen at random to avoid any possible dissection processing effect that would correlate with individual ancestry.

### Total RNA Isolation

RNA was extracted and isolated from each of the three sample replicates from each placenta using TRIZOL reagent (Invitrogen) following manufacturer recommendations. Total RNA was purified using the Qiagen RNAeasy minElute Cleanup kits and RNA quality was determined using Agilent 6000 Nano kits and an Agilent Bioanalyzer.

### Construction of indexed RNA-Seq Libraries

Based on the RNA integrity numbers (RIN values) the two best sample replicates from each placenta were chosen to construct indexed Illumina RNA-Seq libraries. The indices are sample-specific and allow for the pooling and sequencing of all libraries together, thereby minimizing lane and run effects on the RNA-Seq data. Library construction followed a merging of Illumina’s RNA-Seq library preparation and an indexing protocol [53] which introduces barcodes for each library during an enrichment PCR step. Library construction included the following steps: two rounds of mRNA capture with oligo dT magnetic beads, mRNA fragmentation, 1^st^ strand synthesis, 2^nd^ strand synthesis, end repair, index adapter ligation, adapter fill-in, size-selection, indexing/enrichment PCR, and finally quantification. All steps, including SPRI bead reaction clean-ups, were processed in parallel in a 96 well plate, where all samples were randomized across the plate, thereby eliminating any library processing batch effect.

### Sequencing, Base Calling and Mapping

The 80 indexed libraries were pooled in equimolar ratios and sequenced on nine lanes over three runs on the Illumina Genome Analyzer IIx platform. Eight lanes were single-end 76bp (base pair) reads and a ninth lane was a 76bp paired-end run. Base calling was done using Ibis [53] and mapping was performed with TopHat2 [54], a spliced-read mapper which is built on top of the Bowtie mapper [55]. Reads were mapped to the human reference genome build hg19 (GRCh37). Reads were annotated to known Ensembl 70 genes. All count data was normalized (variance stabilized) using protocols described in the DESeq2 package [30]. In instances where data for individuals are used (such as in PCAs), the raw count data from each replicate for each individual were summed and data for individuals was independently normalized with the aforementioned method.

### Apportioning Expression Variation

We decomposed variation in expression level into multiple factors of interest using models derived from those previously established [17, 56]. Data were fit to both a normal and a negative binomial distribution (glm.nb in R), with significant correlations among model estimates (r = 0.9998, p < 2.2e-16). We therefore present all subsequent analyses assuming a normal distribution. Apportionment estimates were calculated using two different components of the data, (1) the sums of squares (SS) and (2) the additive components of variances (σ^2^). The latter is derived from the expected mean squares (EMS) formulas for each explanatory variable [56]. There are several reasons for using these two different parameterizations of the apportionment of expression variance. First, the sum of squares based parameters can be directly compared across ANOVA model types (model I, model II and mixed models). Second, the sum of squares based parameters are more dynamic in that they preclude the possibility of having 0 values. Third, the parameters based on the additive components of variances are a previously published set of parameters that are direct analogs to Wright’s F-statistics (Fst and Fis), a desirable feature that will allow for comparisons between genetic and phenotypic variation [39]. Finally, when using generalized linear models, such as when fitting a negative binomial distribution to the data, the deviance estimates can be used as sums of squares to derive both parameter types. Our simple model for each gene is a model II nested hierarchical ANOVA:

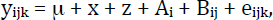

where y is normalized gene expression for the k^th^ sample replicate in the j^th^ individual in the i^th^ group, x and z are technical explanatory variables (x is the number of mapped reads for each library and z the RIN value), and μ is mean expression for any gene g. The group (A), individual (B) and sample replicate (e) effects are assumed to be random with variance σ_A_^2^, σ_B_^2^ and σ^2^, respectively. After removing the variance from technical factors (x and z), the total variance in gene expression can then be apportioned as σ^2^_T_ = σ_A_^2^ + σ_B_^2^ + σ^2^. We summarize the amount of expression variance attributed to groups as σ_A_^2^/σ^2^_T_ and define this correlation coefficient as Mst, the expression variance analog to the standard among-group component of the total genetic variance, Fst [1, 57]. Further we can define the correlation coefficients Met and Mit as the amount of expression variance attributed to sample replicates and error (Met = σ^2^/σ^2^_T_), and to individuals (Mit = σ_B_^2^/σ^2^_T_). Each parameter ranges in value from 0 to 1 and the sum of these parameters, for each gene, equals 1.

Similarly we also estimated a complementary (η^2^) statistic for each explanatory factor, using the sums of squares (SS). In this instance, total gene expression variation can be expressed as SS_T_ = SS_A_ + SS_B_ + SS_e_, and we can subsequently define the parameters Net (SS/SS_T_), Nit (SS_B_/SS_T_) and Nst (SS_A_/SS_T_), which mirror the aforementioned parameters derived from the additive components of variance (Met, Mit, and Mst, respectively). Additionally, we defined Nis as SS_B_/(SS_B_ + SS_e_) to quantify the amount of inter-individual variation relative to the total inter- and intra-individual variation. Finally, we defined Nig as SS_B_/(SS_A_ + SS_B_) to quantify the amount of inter-individual variation relative to the total inter-individual and intergroup variation. We will refer to these two sets of parameters as the apportionment of variance (using σ^2^) and apportionment of variation (using SS) parameters, respectively. An ANOVA table providing further details of the models can be found in Table S1 and S2 in Additional file 1.

We also derived a more complex model for gene expression variation, which accounts for other possible factors that might influence the expression of each gene, namely: sex of the child (s), birth weight of the child (w), birth length of the child (l), manner of birth (c; cesarean or natural), maternal age (f), maternal body mass index (o), if the mother drinks alcohol on a regular basis (d), and if the mother is a vegetarian (v). This is a partially nested model II anova with no interaction:

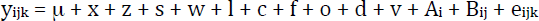

Total variation in expression was apportioned using the η^2^ statistic in a manner similar to that described above, except in this instance all explanatory variables were used and yield:

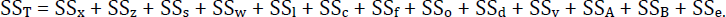

### Mode of Selection Permutations

To determine if variation at each gene may be consistent with a particular mode of selection, a series of permutations were performed, building on the models of Whitehead and Crawford [38]. There are four types of selection to consider: directional, balancing, stabilizing and diversifying. First, directional selection, or simply differential expression (DE), is typified by large variation among groups. To test for directional selection, we permuted individuals among groups 1000 times, maintaining replicate associations, randomly sampled 100 genes for each permutation, and apportioned variation as described above. The 99^th^ percentile of the permuted Nst distribution was taken as a cutoff for extreme Nst values and thus DE genes.

The second type of selection is balancing selection, typified by high among individual variation along with low among population variation [38, 58, 59]. Balancing selection was examined by permuting sample replicates among individuals within groups 1000 times (randomizing inter-individual differences), randomly sampling 100 genes for each permutation and apportioning variation. The parameter Nig was used to identify genes with significantly more variation among individuals than among groups. The 99^th^ percentile of the permuted Nig distribution was taken as a cutoff for extreme Nig values.

The other types of selection are stabilizing selection (characterized by low among individual variation) and diversifying selection (characterized by high individual variation). In these later two modes of selection we specifically assume that selection does not vary spatially and is thus uniform across populations. To identify profiles consistent with stabilizing or diversifying selection, we generated a random distribution of inter-individual variances as follows. Gene expression was normalized across all genes, so that all genes have the same mean expression. We then randomly selected the expression level of any one gene from each individual to create a new artificial gene. We did this 10,000 times and calculated the variance across all individuals with no regard for population association. The 1^st^ percentile and 99^th^ percentile of this distribution were taken as cutoffs for stabilizing and diversifying selection, respectively.

### GO & KEGG Enrichment

Enrichment in GO (Gene Ontology) categories and KEGG (Kyoto Encyclopedia of Genes and Genomes) pathways were performed using the GOSeq [60] R package, designed to account for read count biases in transcript length from RNA-Seq data. In all enrichment analyses we present results using two false discovery rate (FDR) cutoffs - a high confidence FDR of 5% and a moderate confidence FDR of 20%. All p-values and FDRs are provided in supplementary materials.

### Coexpression Modules, Network Construction and Profile Partitioning

Gene coexpression modules were identified using a weighted gene coexpression network analysis (WGCNA) [61]. Network graphs were constructed using the graph.adjacency function from the igraph package. Interacting genes, used to build the network, were identified by using the dissimilarity values from the WGCNA analysis but limiting them to those that were additionally significant in a linear regression correlation analysis at an FDR of 1%. For our data and this analysis, an FDR of 1% corresponds roughly to a Pearson’s r >= 0.6 and a dissimilarity value <=0.3. Gene expression profile partitioning was performed using k-means clustering (kmeans function in the stats package in R). Data from individuals was used and expression at each gene was normalized, prior to clustering, to have a mean of 0 and a standard deviation of 1.

### Statistical Analyses

All analyses were carried out in the programming language R [62] with in house scripts and the aforementioned packages. All p-values are Benjamini-Hochberg [63] adjusted using the p.adjust function from the R package stats, and significance is taken at a p-adjust/FDR of 0.05, unless stated otherwise.

### Validation

We validate three genes with extreme Mst values by performing rt-qPCR. In this analysis one of the two original sample replicates was used, along with the third RNA sample processed at the same time as the study samples but not used to create an RNA-Seq library. The Maxima SYBR Green qPCR Master Mix from Fermentas was used following the manufacturer’s instructions. Primer sequences are presented in Table S3 in Additional file 1.

## Additional Material

Additional file 1 (word doc; .doc), Supplementary Table and Figures: contains all supplementary figures and tables

Additional file 2 (excel doc, .xls), PCA Enrichment: contains GO and KEGG enrichment results for the top 4 principle components derived from total gene expression variation across individuals.

Additional file 3 (excel doc, .xls), Selection Mode Enrichment: contains GO and KEGG enrichment results for genes under different forms of selection and for particular co-expression modules. Results are partitioned onto different worksheet/tabs as denoted in the text.

Additional file 4 (excel doc, .xls), Trait Enrichment: contains GO and KEGG enrichment results for those genes associated with each technical, biological or dietary factor modeled in or full model of gene expression variation.

Additional file 5 (excel doc, .xls), Gene Selection Categories: contains four lists of Ensembl identifiers for genes classified as being influenced by Directional, Stabilizing, Diversifying or Balancing selection.

## Author Contributions

MS conceived the project. DAH, CSM and GLF collected the samples. DAH performed the laboratory analyses and DAH, MK, and ZH analyzed the data. DAH and MS wrote the manuscript. PK and GS facilitated and advised analyses.

## Acknowledgments

We would like to thank all donors for their participation, and the nursing staff at Northside Hospital in Atlanta, Georgia for their assistance in the sample collection. The Max Planck Society and a grant for visiting young scientists from the Chinese Academy of Science (2009YB1-12) funded the project. The funders had no role in study design, data collection and analysis, decision to publish, or preparation of the manuscript.

